# NF90 Interacts with Components of RISC and Modulates Association of Ago2 with mRNA

**DOI:** 10.1101/2021.09.23.461467

**Authors:** Giuseppa Grasso, Charbel Akkawi, Celine Franckhauser, Rima Nait-Saidi, Maxime Bello, Jérôme Barbier, Rosemary Kiernan

**Affiliations:** UMR9002 CNRS-UM, Institut de Génétique Humaine-Université de Montpellier, Gene Regulation lab, Montpellier, 34396, France

**Author notes:** contributed equally (Giuseppa Grasso, Charbel Akkawi).

## Abstract

Nuclear Factor 90 (NF90) is a double-stranded RNA-binding protein involved in a multitude of different cellular mechanisms such as transcription, translation, viral infection and mRNA stability. Recent data suggest that NF90 might influence the abundance of target mRNAs in the cytoplasm through miRNA- and Argonaute 2 (Ago2)-dependent activity. Here, we identified the interactome of NF90 in the cytoplasm, which revealed several components of the RNA-induced silencing complex (RISC) and associated factors. Co-immunoprecipitation analysis confirmed the interaction of NF90 with the RISC-associated RNA helicase, Moloney leukemia virus 10 (MOV10), and other proteins involved in RISC-mediated silencing, including Ago2. Furthermore, NF90 association with MOV10 and Ago2 was found to be RNA-dependent. Glycerol gradient sedimentation of NF90 immune complexes indicates that these proteins occur in the same protein complex. At target RNAs predicted to bind both NF90 and MOV10 in their 3’ UTRs, NF90 association was increased upon loss of MOV10 and *vice versa*, suggesting that the two proteins may compete for the binding of common target mRNAs. Interestingly, loss of NF90 led to an increase in association of Ago2 as well as a decrease in the abundance of the target mRNA. Similarly, during hypoxia, the binding of Ago2 to vascular endothelial growth factor (VEGF) mRNA increased after loss of NF90, while the level of VEGF mRNA decreased. These findings suggest a role for NF90 in the regulation of RISC-mediated silencing which stabilizes target mRNAs, such as VEGF, during cancer-induced hypoxia.

## INTRODUCTION

Nuclear Factor 90 (NF90) is a double-stranded RNA-binding protein (RBP) that is involved in a plethora of different cellular processes and pathways, such as transcription, splicing, translation and mRNA stability or degradation [1]. More recently, NF90 was also linked to microRNA (miRNA) biogenesis and circular RNA (circRNA) stability [2, 3]. NF90 is a ubiquitous and generally abundant protein that has been shown to shuttle from the nucleus to the cytoplasm depending on its phosphorylation status and as a result of several stimuli, such as viral infection or hypoxia [4–6]. During viral infection, cytoplasmic NF90 can bind viral RNAs to enhance or inhibit viral replication, depending on the type of virus [7]. Besides viral RNAs, NF90 can associate with cellular mRNAs to increase their stability or influence their translation [8, 9]. For instance, NF90 was recently shown to play a role in mitosis by competing with Staufen-mediated mRNA decay (SMD) for the binding of mitotic mRNAs [10]. NF90 contains two tandem double-stranded RNA-binding motifs (dsRBMs) that were shown to contribute to the binding of the same RNA molecule simultaneously [11, 12]. It was shown that NF90 is able to recognize specific RNAs structures, such as minihelix, and that this structure is sufficient for NF90 binding [13, 14]. A sequence motif has also been shown to positively influence NF90 binding to short mRNAs [11]. Therefore, the precise RNA binding mode of NF90 is still not clear and may depend on the type of RNA. Nevertheless, NF90 RNA binding activity is strongly influenced by the heterodimerization with its protein partner Nuclear Factor 45 (NF45) which leads to thermodynamic stabilization of the complex and enhanced affinity for RNA substrate [15].

Recent findings implicate NF90 in mRNA stability and translation through miRNAs [16], which could suggest an involvement in RISC-mediated gene silencing. RISC-mediated gene silencing is a well-known post-transcriptional gene regulation mechanism that, in humans, was shown to promote translational inhibition and degradation of target mRNAs mainly by recruiting Ago2 protein [17]. Ago2, guided by the sequence complementarity of a miRNA, is able to diffuse along the 3’ UTR of target mRNAs recognizing miRNA recognition elements (MRE) and recruiting effector proteins, such as deadenylases and 5’-to-3’ exonucleases, in order to inhibit translation and/or trigger mRNA degradation [18]. Interestingly, characterization of the Ago2 interactome identified NF90/NF45 heterodimer as an interactant of Ago2 in the cytoplasm [19], which could suggest a role for NF90/NF45 in RISC-mediated activities.

For Ago2 lateral diffusion to be efficient, it needs the activity of a helicase to disrupt occlusive secondary RNA structures that could interfere with its binding. Moloney leukemia virus 10 (MOV10) is an ATP-dependent helicase that belongs to the Up frameshift (UPF)-like helicase superfamily 1 (SF1). MOV10 binds ssRNA and translocates 5’-to-3’ along the target RNA [20]. It was initially shown to inhibit Human Immunodeficiency Virus type 1 (HIV-1) and Hepatitis C virus replication as well as LINE-1 retrotransposition [21, 22]. In addition, MOV10 was found to co-localize in P-bodies together with Ago2 and other factors involved in RISC, identifying a role for MOV10 in miRNA-mediated regulation [23]. MOV10 binds in close proximity to UPF1 binding sites to resolve structures and displace RBPs from the 3’ UTR of the target mRNA, thereby exposing the MRE for Ago2 binding [20].

Within a mRNA target, MOV10 frequently binds to the 3’ UTR, specifically at regions with low conservation and upstream of local secondary structures, consistent with its 5’-to-3’ directional unwinding activity [20]. Moreover, for efficient loading and unwinding, MOV10, like many RNA helicases, needs a single-stranded region of RNA adjacent to a duplex [24]. Consistent with its role in RISC-mediated silencing, MOV10 can regulate the abundance of the mRNAs to which it binds. In particular, it was shown that depletion of MOV10 inhibits translational suppression leading to global stabilization of its target mRNAs [20]. However, in contrast with this observation, MOV10 was also found to increase the expression of a limited subset of mRNAs, in the presence of fragile X mental retardation protein 1 (FMRP1), by inhibiting Ago2 binding. Therefore, the concomitant binding of MOV10 and FMRP on the same mRNA can inhibit the canonical role of MOV10 in RISC [25, 26].

Here, we show that cytoplasmic NF90 interacts with proteins involved in translational repression, RNA stability, degradation and viral replication. We determined that NF90 interacts with MOV10 as well as Ago2 in an RNA-dependent fashion. Using published CLIP data of MOV10 and NF90, we showed that both proteins can bind the same target mRNAs using RNA immunoprecipitation (RIP) analysis. Upon loss of MOV10, association of NF90 with the mRNA targets increased. Similarly, after loss of NF90/NF45, we detected an increase in the association of MOV10 to the selected target mRNAs. To determine whether NF90 binding might impact association of RISC with the mRNA targets, we performed RIP analysis for Ago2. Association of Ago2 with target mRNAs increased following loss of NF90 and the abundance of the target mRNAs was concomitantly decreased. Hypoxia leads to stabilization of specific NF90-associated mRNAs, such as vascular epithelial growth factor (VEGF) mRNA. Interestingly, during hypoxia, the abundance of VEGF mRNA decreased after loss of NF90/NF45, while its association with Ago2 significantly increased. These data suggest that NF90 may be involved in RISC-mediated gene silencing by regulating MOV10 and Ago2 association with target mRNAs. Moreover, our results suggest that this novel role of NF90 might be implicated in the response to cancer-induced hypoxia.

## MATERIAL AND METHODS

### Cell culture, stable cell line production and cellular treatments

Human HEK293T cell line was grown in Dulbecco’s Modified Eagle’s Medium containing high glucose with HEPES modification (Sigma-Aldrich®, D6171), supplemented with 10% fetal bovine serum (PAN Biotech, 8500-P131704), 1% penicillin-streptomycin (v/v) (Sigma Aldrich^®^, P4333) and 1% L-glutamine (v/v) (Sigma Aldrich®, G7513). Cells were cultured at 37°C in a humidified atmosphere containing 5% CO_2_. For experiments using RNAi, 3 × 10^6^ cells were seeded in 100 mm culture dishes at the day of siRNA transfection or at 5 × 10^6^ cells for experiments using RNAi together with CoCl_2_ treatment.

Plasmid encoding pOZ-NF90-FLAG-HA (pOZ-NF90-FH) were cloned using pOZ-N-FH vector, as previously described [27]. Briefly, lentiviral particles expressing NF90 were produced in HEK293T cells by transfecting plasmids using calcium-phosphate. HEK293T were transduced using Polybrene infection/transfection reagent (Sigma-Aldrich^®^, TR-1003), according to the manufacturer’s instructions. After 7 days, selection of transduced cells was carried out by magnetic affinity sorting with antibody against IL2 to achieve a pure population. HEK293T stably expressing NF90 (pOZ-NF90-Flag-HA HEK293T) were grown in the same conditions as HEK293T. pOZ-NF90-FH HEK293T were seeded at 1&10^7^ in 150 mm culture dishes the day prior to protein extraction. For hypoxia treatment, HEK293T were treated for 24 h with 500 µM of CoCl_2_ (Sigma, 15862-1ML-F) or the same volume of water as a control, where indicated.

### Transfection of small interfering RNAs

Double-stranded RNA oligonucleotides used for RNAi were purchased from Eurofins MWG Operon or Integrated DNA Technologies. Sequences of small interfering RNAs (siRNAs) used in this study have been described previously [13] and are shown in Supplementary Table S1. HEK293T cells were transfected with siRNA (30 nM final concentration) using INTERFERin® siRNA transfection reagent (PolyPlus Transfection) according to the manufacturer’s instructions. The transfection was carried out the day of seeding and cells were collected for protein extraction approximately 65 h after transfection.

### Immunoblot

HEK293T and pOZ-NF90-FH HEK293T were lysed using RIPA buffer (50 mM Tris-HCl pH=7.5, 150 mM NaCl, 1 % NP40, 0.5 % Sodium Deoxycholate, 0.1 % SDS, Halt™ Phosphatase Inhibitor Cocktail (Thermo Fisher Scientific)), unless otherwise indicated. Protein extracts were immunoblotted using the indicated primary antibodies (Table S2) and anti-mouse, anti-rabbit or anti-rat IgG-linked HRP secondary antibodies (GE Healthcare) followed by ECL (Advansta).

### Cytoplasmic extracts and co-immunoprecipitation analysis

HEK293T were seeded in 150 mm culture dishes the day prior to protein extraction. Cytoplasmic proteins were extracted using a mild lysis buffer (10 mM Hepes pH 7.9, 10 mM KCl, 0.1 mM EDTA pH 8.0, 2 mM MgCl2, 1 mM DTT, EDTA-free protease and phosphatase inhibitor). The cell pellet was incubated for 10 min on ice, adding 0.07% NP-40 and incubating for additional 10 min on ice. After centrifugation, 1 mg of lysates were incubated at 4°C overnight with 2 μg of antibodies recognizing NF90, Ago2 and IgG controls and protein A/G PLUS-Agarose beads (Santa Cruz, sc-2003). Beads were then washed twice with IP buffer (150 mM KCl, 20 mM Tris pH 7.5, 0.05% NP-40, 0.1% Tween, 10% glycerol, 5 mM MgCl_2_, 1 mM DTT and EDTA-free protease and phosphatase inhibitor). Samples were treated with RNAse A/T1 mix (ThermoFisher Scientific, EN0551) for 30 min at room temperature, incubating on a rotating wheel. After incubation, beads were washed three times with IP buffer as aforementioned and 2X Laemmli buffer was added directly to the beads.

### Tandem Immunoprecipitation and mass spectrometry

For mass spectrometry, cytoplasmic extracts were obtained as aforementioned. Tandem immunoprecipitation (Flag and HA) was carried out using 10 mg of cytoplasmic extract. Flag IP was performed using EZview™ Red ANTI-Flag^®^ M2 Affinity gel (SigmaAldrich, F2426), following the manufacturer’s instructions. Washes were carried out 3 times as aforementioned and protein complexes were eluted by competition performing 2 consecutive elutions using Flag elution buffer (250 ng/µl FLAG^®^ Peptide (SigmaAldrich, F3290), diluted in IP buffer) incubating for 1 h at 4°C on a rotating wheel. HA IP was performed using the elutions obtained from the first IP incubated with Pierce™ Anti-HA Agarose beads (ThermoScientific, 26181) for 2 h at 4°C on a rotating wheel. Washes were carried out 5 times as aforementioned and elutions were performed using HA elution buffer (400 ng/ µl HA peptide (ThermoScientific, 26184), diluted in IP buffer) for 1 h at 4°C on a rotating wheel. Following elution, beads were removed using Pierce™ Centrifuge Columns (ThermoScientific, 11894131), as specified by manufacturer’s instructions. Silver-staining was performed according to the manufacturer’s instructions (Silverquest, Invitrogen). Mass spectrometry was performed at Taplin facility, Harvard University, Boston, MA.

### Glycerol gradient sedimentation

Glycerol gradient sedimentation was performed as described previously [27]. Briefly, NF90-associated proteins were purified by performing FLAG IP on pOZ-NF90-FH HEK293T cytoplasmic fraction, as aforementioned. One ml layers of glycerol (final concentration 15 to 35%–20 mM Tris pH 7.5, 0.15 M KCl, 2.5 mM MgCl2, 0.05% NP-40, 0.1% Tween) were layered into centrifugation tubes (13 × 51 mm Ultra-Clear Tubes, Beckman). A linear gradient was obtained after 12 h of diffusion at 4°C. Flag elution from pOZ-NF90-FH HEK293T immunoprecipitation was loaded on top of the glycerol gradient. Complexes were fractionated by ultracentrifugation in an SW 55Ti rotor (Beckman) at 30,000 rpm for 18 h at 4°C. 25 fractions of 200 μL were collected from top of the gradient. An equal volume of fractions was resolved by SDS-PAGE and immunoblotted with indicated antibodies.

### RNA immunoprecipitation (RIP)

RIP was performed as previously described [13]. Briefly, HEK293T were seeded in 100 mm culture dishes and transfected with siRNAs the day of seeding or treated with CoCl_2_, as aforementioned. Cells were harvested ∼ 65 h after the siRNA treatment or 24 h after CoCl_2_ treatment and lysed for 10 min in RIP buffer (20 mM HEPES, pH 7.5, 150 mM NaCl, 2.5 mM MgCl_2_•6H_2_O, 250 mM sucrose, 0.05% (v/v) NP-40 and 0.5% (v/v) Triton X-100) containing 20 U ml^−1^ of RNasin (Promega), 1 mM DTT, 0.1 mM PMSF and EDTA-free protease and phosphatase inhibitor. After centrifugation, lysates were incubated for 4 h at 4°C with 2 μg of antibodies recognizing NF90, MOV10, Ago2 and IgG control and then incubated for 1 h at 4°C with Dynabeads™ Protein A or G (ThermoFisher Scientific). After incubation, beads were washed five times with RIP buffer for 5 min at 4°C and RNA was extracted using TRIzol (Thermo Fisher Scientific) according to the manufacturer’s instructions. RNA was treated with DNAse I (Promega) and RT was performed using SuperScript™ III Reverse Transcriptase (ThermoFisher Scientific) according to the manufacturer’s instructions. cDNA was treated with RNAse H (ThermoFisher Scientific) and the samples were used to perform qPCRs using LightCycler™ 480 SYBR Green I Master mix (Roche), according to the manufacturer’s instructions, using the primers shown in Supplementary Table S3.

### Bioinformatic analyses

Enhanced UV crosslinking followed by immunoprecipitation (eCLIP) data for NF90 were obtained from Nussbacher and Yeo [28] and retrieved from the NCBI database (NF90 eCLIP: ENCSR786TSC). Individual-nucleotide-resolution UV crosslinking and (iCLIP) data for MOV10 were obtained and retrieved from the NCBI database (MOV10 iCLIP: GSE51443). MOV10 iCLIP data was lifted to hg38 genome annotation using UCSC liftOver tool. Peaks were filtered based on Fold Change (FC ≥ 1.5) and p-value (Bonferroni-Adj P-val ≤ 0.05). Bigwig files from different replicates were merged using bigWigMerge v2. Statistical analyses were performed using RStudio v1.4. Gene ontology was performed using DAVID Functional Annotation Tool database version 6.8 (https://david.ncifcrf.gov).

## RESULTS

### NF90 interacts with proteins involved in translational repression and RNA processing

In order to understand the role of NF90 in the cytoplasm, we determined its cytoplasmic interactome by performing mass spectrometry on a HEK293T cell line stably overexpressing NF90-FLAG-HA (Figure S1A), after tandem affinity purification of the cytoplasmic protein fraction (Figure S1B). Excluding the proteins for which one peptide or more was found in the mock HEK293T sample, 318 proteins were detected associated with NF90 in the cytoplasm (Table S4). As expected, the most abundant protein identified was NF90. ILF2 (NF45), a well-known NF90 protein partner, was found among the most abundant interactants. Interestingly, several proteins associated with RISC were identified among the interactants. Notably, the 5’-to-3’ helicase MOV10, was highly represented. Ago2, the main component of RISC, was also associated with NF90, as reported previously [19]. Gene ontology of the significantly enriched NF90 interactants identified a number of pathways, such as translation, translational initiation and its regulation, mRNA processing and regulation of mRNA stability (Figure 1A). These findings are consistent with previous observations suggesting the implication of NF90 in the regulation of mRNA stability and mRNA translation for specific target mRNAs [6, 16].

**Figure 1.**
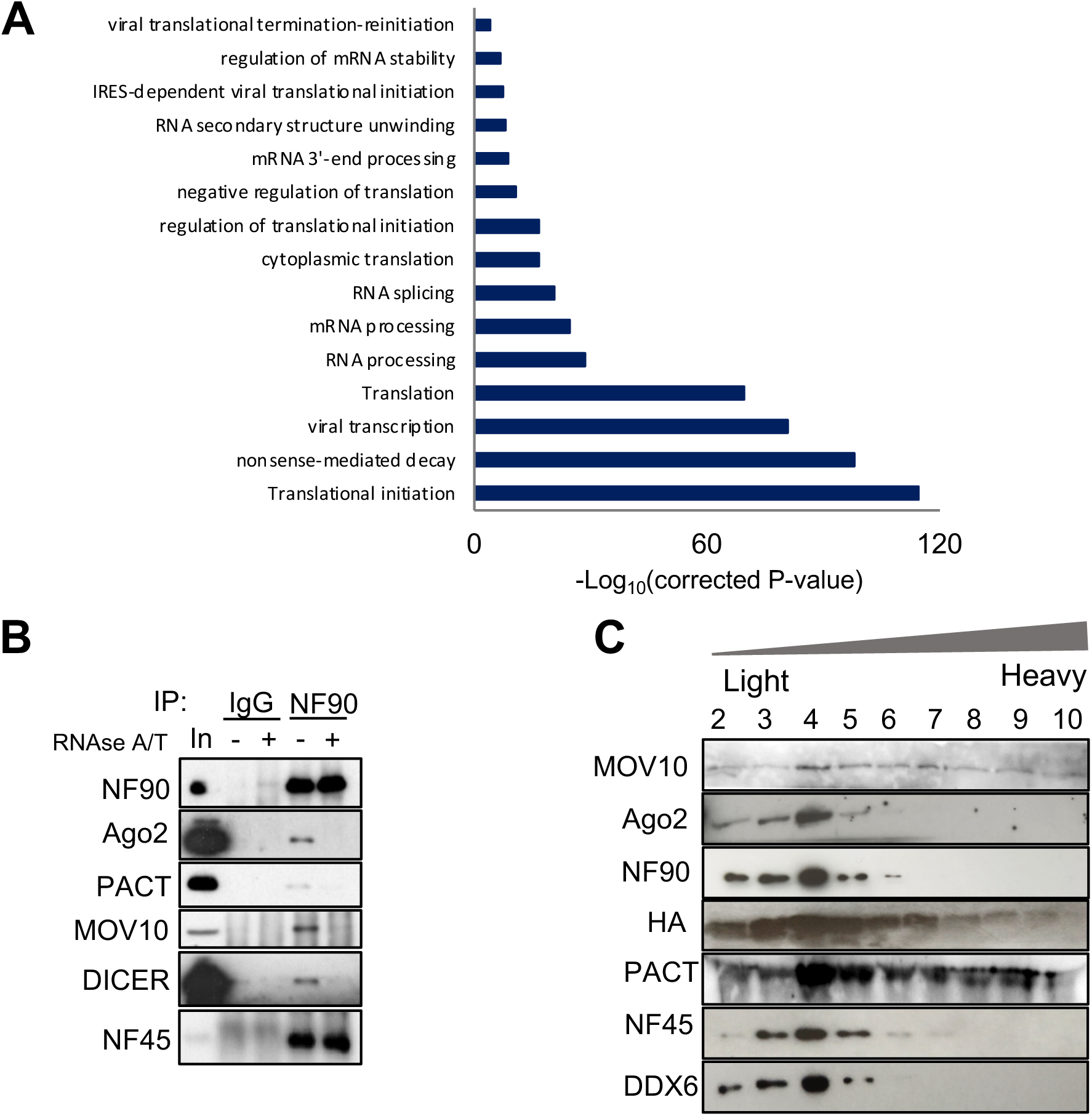
NF90 interacts with proteins involved in translational repression and RNA processing. **(A)** Gene ontology analyses (cellular pathways) of NF90-associated proteins identified by mass spectrometry. **(B)** Cytoplasmic extracts from HEK-293T cells were used for immunoprecipitation using an anti-IgG control or anti-NF90 antibody followed by RNAse A/T treatment or mock treatment, as indicated. Immunoprecipitates and an aliquot of extract (In) were analyzed by western blot, using the indicated antibodies. **(C)** FLAG immunoprecipitates of cytoplasmic extracts of NF90-FH overexpressing HEK293T cells were separated by glycerol gradient sedimentation. NF90-containing fractions were analyzed by western blot using the antibodies indicated.

Since NF90 interactants are well-known RNA-binding proteins (Supplementary Figure S1C), we wondered if their interaction with NF90 was RNA dependent. In order to validate the results of mass spectrometry using endogenous proteins and to determine if the binding of NF90 is RNA dependent, we performed co-IP in HEK293T cell line, with and without RNAse A/T treatment. The results confirmed the binding of endogenous NF90 to PACT, MOV10 and Dicer and furthermore showed that these interactions are RNA-dependent (Figure 1B). On the other hand, the binding of NF90 to NF45 was RNA-independent (Figure 1B), as previously reported [29, 30]. Although the binding of Ago2 to NF90 was previously reported to be RNA-independent [19], under the conditions used in this study, NF90 binding to Ago2 was RNA-dependent, in agreement with a recent study [16]. These findings identify an RNA-mediated interaction between NF90 and RISC-silencing complex in the cytoplasm.

To determine whether RISC subunits can be found in the same complex with NF90, we performed glycerol gradient sedimentation of immunopurified NF90-Flag-HA complexes (Figure S1D and S1E). NF90 was detected in the top 10 fractions (data not shown) where it co-sedimented with MOV10, AGO2, PACT, DDX6 and NF45 (Figure 1C). In agreement with these results, NF90 has been shown to co-sediment with Ago2, NF45 and other helicases, such as DDX47, DDX36 and DDX30, on a sucrose gradient [19]. These findings suggest that cytoplasmic NF90/NF45 co-sediments with subunits of RISC.

### NF90 and MOV10 can bind the same target mRNAs

Among the most abundant interactants of NF90 was MOV10 helicase that is required for optimal RISC activity. To further investigate the interaction between NF90 and MOV10 and a possible role in RISC function, we took advantage of an enhanced UV crosslinking followed by immunoprecipitation of NF90 (eCLIP) [28] and an individual-nucleotide resolution UV crosslinking and IP of MOV10 (iCLIP) [26] datasets. Analysis of MOV10 iCLIP identified 1103 mRNAs significantly bound by MOV10. Approximately half of the bound mRNAs (542 mRNAs) contained a MOV10 peak in the 3’UTR (Figure 2A), consistent with previous findings suggesting that MOV10 mainly binds 3’UTRs of mRNAs [20]. The remaining MOV10-bound mRNAs contained peaks in introns (279 mRNAs), exons (255 mRNAs) and 5’ UTRs (27 mRNAs) (Figure 2A). On the other hand, analyses of NF90 eCLIP suggest that the majority of mRNAs significantly associated with NF90 were bound in their introns (3217 out of 3942 NF90-bound mRNAs). However, some mRNAs were also bound in their in exons, 5’ UTRs and 3’ UTRs (301, 70 and 354, respectively) (Figure 2B).

**Figure 2.**
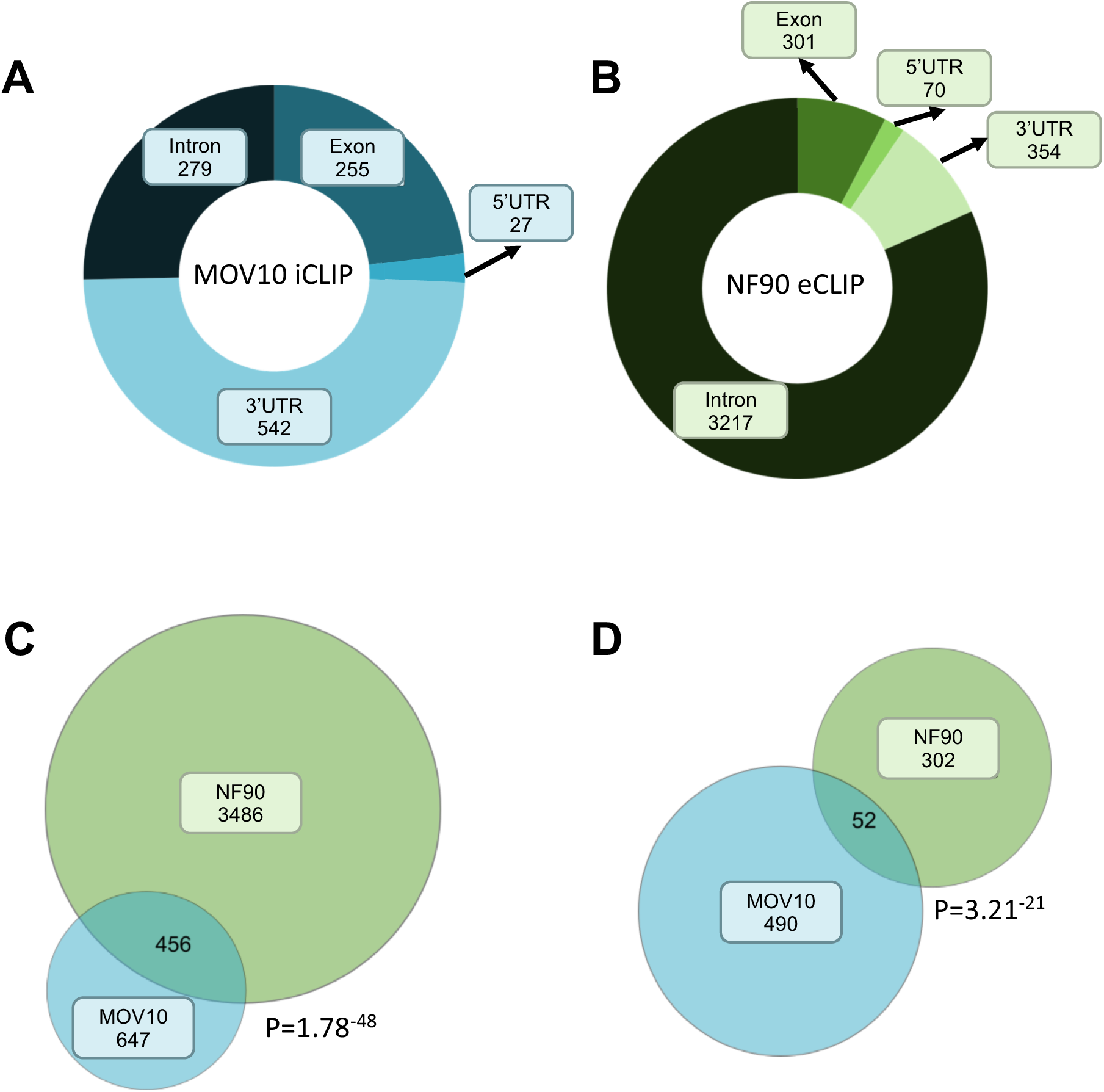
NF90 and MOV10 can bind common target mRNAs. **(A)** MOV10 iCLIP data were analyzed to show the distribution of MOV10 binding along MOV10-associated mRNAs. **(B)** NF90 eCLIP data were analyzed to show the distribution of NF90 binding along NF90-associated mRNAs. **(C)** Venn diagram showing the intersection between target RNAs bound by MOV10 or NF90. P-value was obtained using Fisher’s exact test. **(D)** Venn diagram showing the intersection between target RNAs bound in the 3’ UTR by MOV10 or NF90. P-value was obtained using Fisher’s exact test.

Since the interaction between NF90 and MOV10 was found to be RNA-dependent (Figure 1B), we wondered if these two proteins could bind the same mRNAs. Intersection of the mRNAs bearing at least one peak of NF90 and MOV10 identified 456 mRNAs that could be potentially bound by both proteins (Figure 2C). Interestingly, this corresponds to around 41% of all mRNAs bound by MOV10 and it is significantly enriched (P= 1.7e-48, Fisher’s exact test). Next, since the unwinding of 3’ UTRs by MOV10 is implicated in RISC-mediated silencing, we wondered whether NF90 and MOV10 were associated with the 3’UTRs of common mRNAs. We identified 52 mRNAs that bear at least one peak of both NF90 and MOV10 in their 3’ UTRs (Figure 2D), which is significantly enriched (P=3.21e-21, Fisher’s exact test). These findings suggest that NF90 and MOV10 can bind the 3’ UTR of a common set of target mRNAs.

### NF90 and MOV10 influence each other’s binding to target mRNAs

MOV10 has been shown to facilitate RISC-mediated silencing [23] while, on the contrary, NF90 was found to increase specific target mRNAs stability [9]. We therefore wondered if NF90 could interfere with the binding of MOV10 to the target mRNAs and *vice versa*. Therefore, in order to understand the function of NF90 and MOV10 binding to the same target mRNAs, we performed RNA immunoprecipitation (RIP) in HEK293T cell line after RNAi against NF90 and NF45 (NF90/NF45), MOV10 or a non-targeting control (Scr), followed by quantitative PCR (qPCR) of target mRNAs selected on the basis of MOV10 iCLIP and NF90 eCLIP analyses (Figure 2D and Figure S2).

RIP of NF90, MOV10 or IgG control was performed after downregulation of MOV10 (Figure S3A). Loss of MOV10 did not significantly affect the abundance of the target mRNAs (Figure S3B). RIP results revealed that MOV10 association with the selected target mRNAs was significantly decreased after downregulation of MOV10, as expected (Figure 3). On the other hand, NF90 binding to the same target mRNAs was significantly increased after downregulation of MOV10, while its binding to the negative control, H2BC1, did not significantly change (Figure 3).

**Figure 3.**
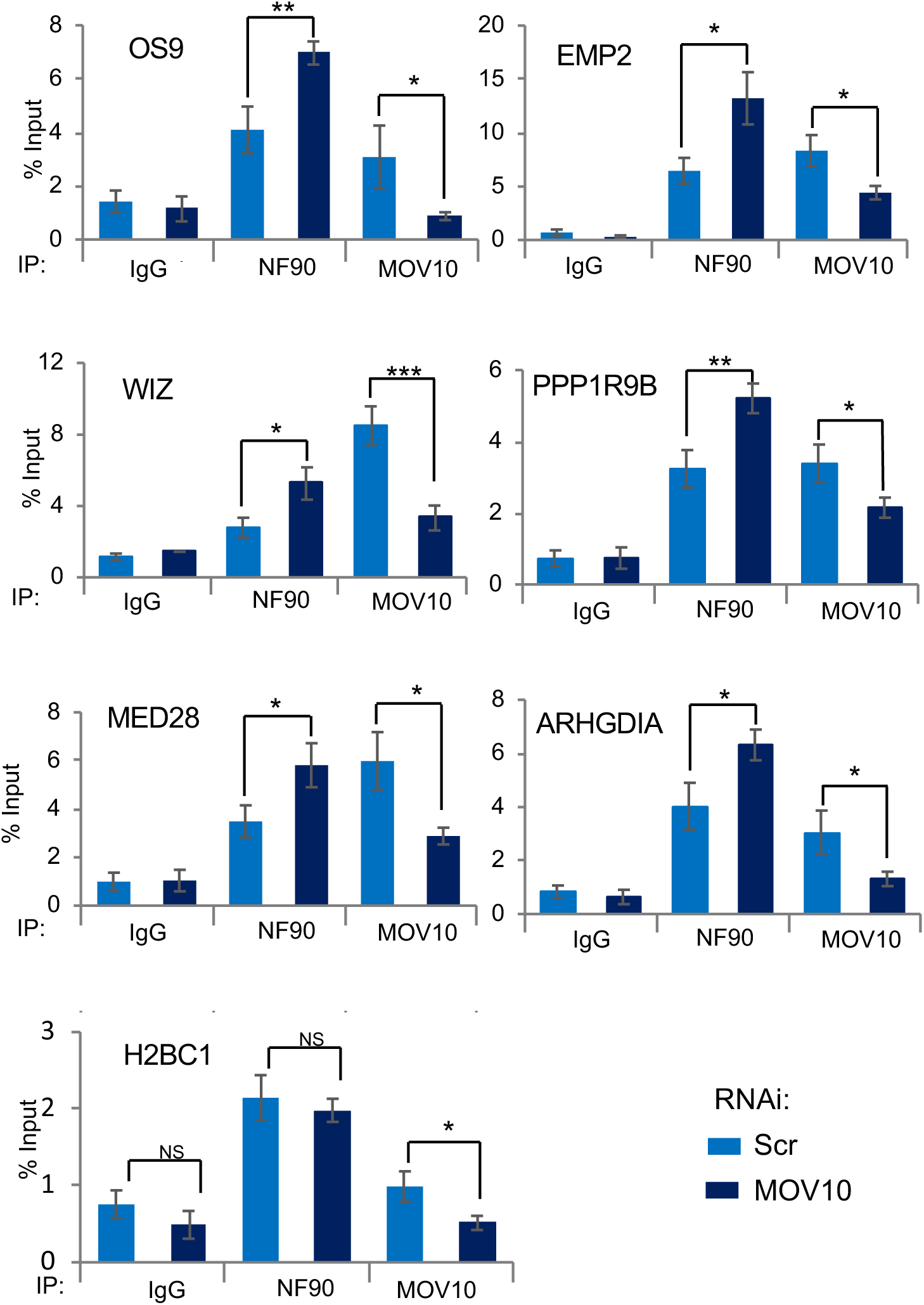
MOV10 modulates NF90 association with common target mRNAs. RIP analysis of HEK293T cells transfected with MOV10-targeting siRNA or a non-targeting control (Scr), as indicated. RIPs were performed using anti-NF90, anti-MOV10 or a control IgG antibody, as indicated. Immunoprecipitates were analyzed using RT-qPCR. Data represent mean ± SEM obtained from 4 independent experiments *(*P < 0.05, **P < 0.01, ***P < 0.001*, independent Student’s *t* test).

In order to further investigate this mechanism, we performed RIP of NF90, MOV10 or IgG control after downregulation of NF90 and its protein partner NF45 (Figure S4A). Interestingly, the loss of NF90/NF45 significantly decreased the total level of the selected target mRNAs (Figure S4B), consistent with its role in increasing mRNA stability [9]. As expected, NF90 association with the target mRNAs was significantly decreased after NF90/NF45 downregulation (Figure 4). On the other hand, RIP analysis revealed that NF90/NF45 downregulation led to a significant increase in MOV10 association with the selected target mRNAs while its binding to the negative control, H2BC1, did not significantly change (Figure 4). These results suggest that the binding of NF90 and MOV10 at common target mRNAs is mutually influenced by the presence of the other factor.

**Figure 4.**
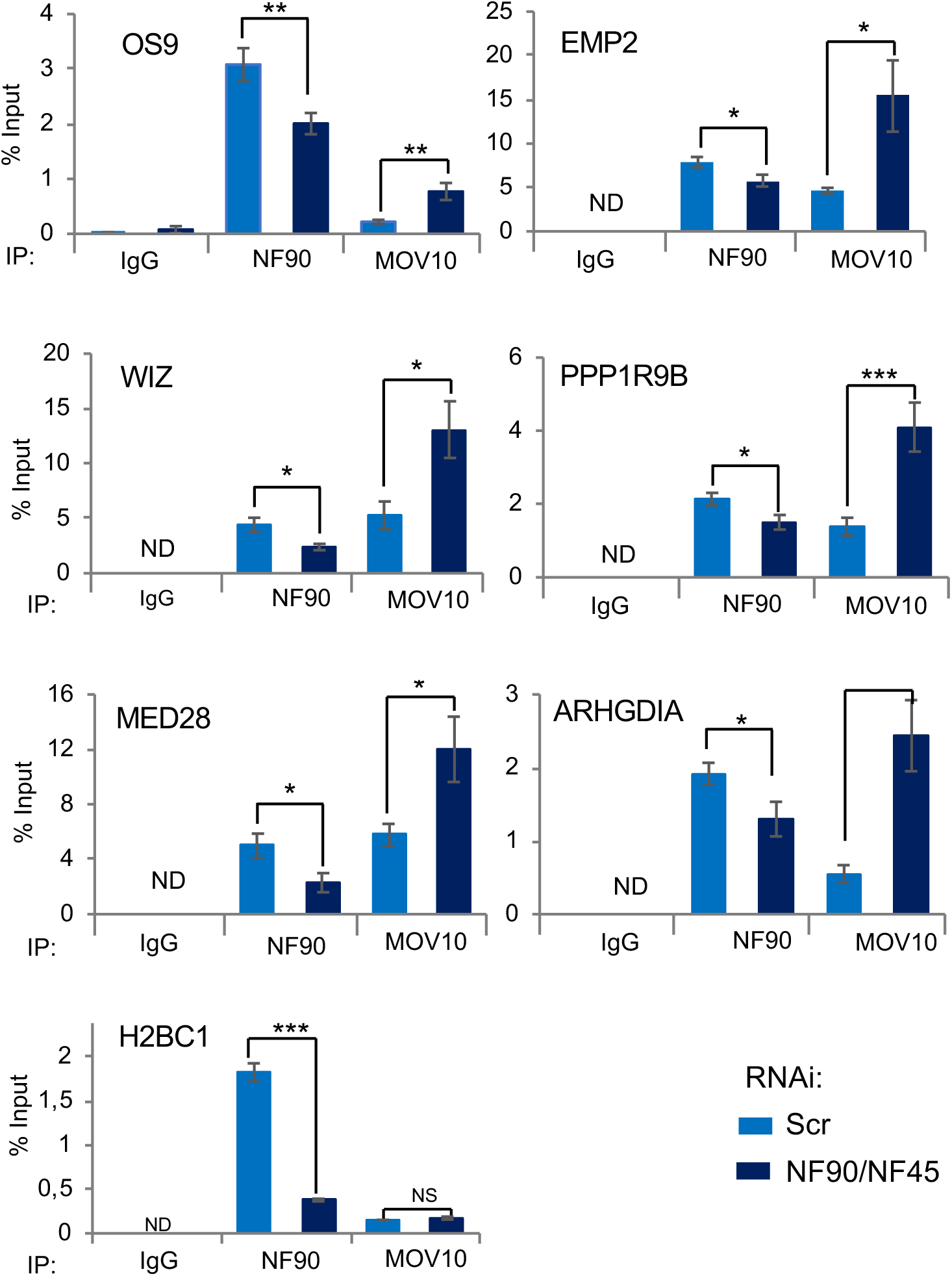
NF90 modulates MOV10 association with common target mRNAs. RIP analysis of HEK293T cells transfected with NF90/NF45-targeting siRNAs or a non-targeting control (Scr), as indicated. RIPs were performed using anti-NF90, anti-MOV10 or a control IgG antibody. Immunoprecipitates were analyzed using RT-qPCR. ND indicates ‘Not Detected’. Data represent mean ± SEM obtained from 4 independent experiments *(*P < 0.05, **P < 0.01, ***P < 0.001*, independent Student’s *t* test).

### Downregulation of NF90/NF45 complex increases Ago2 binding to target mRNAs

MOV10 is known to promote Ago2 association to mRNAs, thereby enhancing RISC-mediated silencing. Since NF90 and MOV10 influence each other’s binding to common target mRNAs, we wondered if alteration of the association NF90/NF45 complex could have similar effects on Ago2 binding to target mRNAs. To this end, we performed RIP of Ago2 or IgG control after downregulation of NF90/NF45. Depletion of NF90/NF45 had no effect on Ago2 expression in extracts or immunoprecipitates (Figure S5A, B). Notably, loss of NF90/NF45 significantly increased Ago2 binding to the target mRNAs tested, while the negative control mRNA, H2BC1, which was poorly associated with Ago2, was not significantly increased relative to the IgG control (Figure 5).

**Figure 5.**
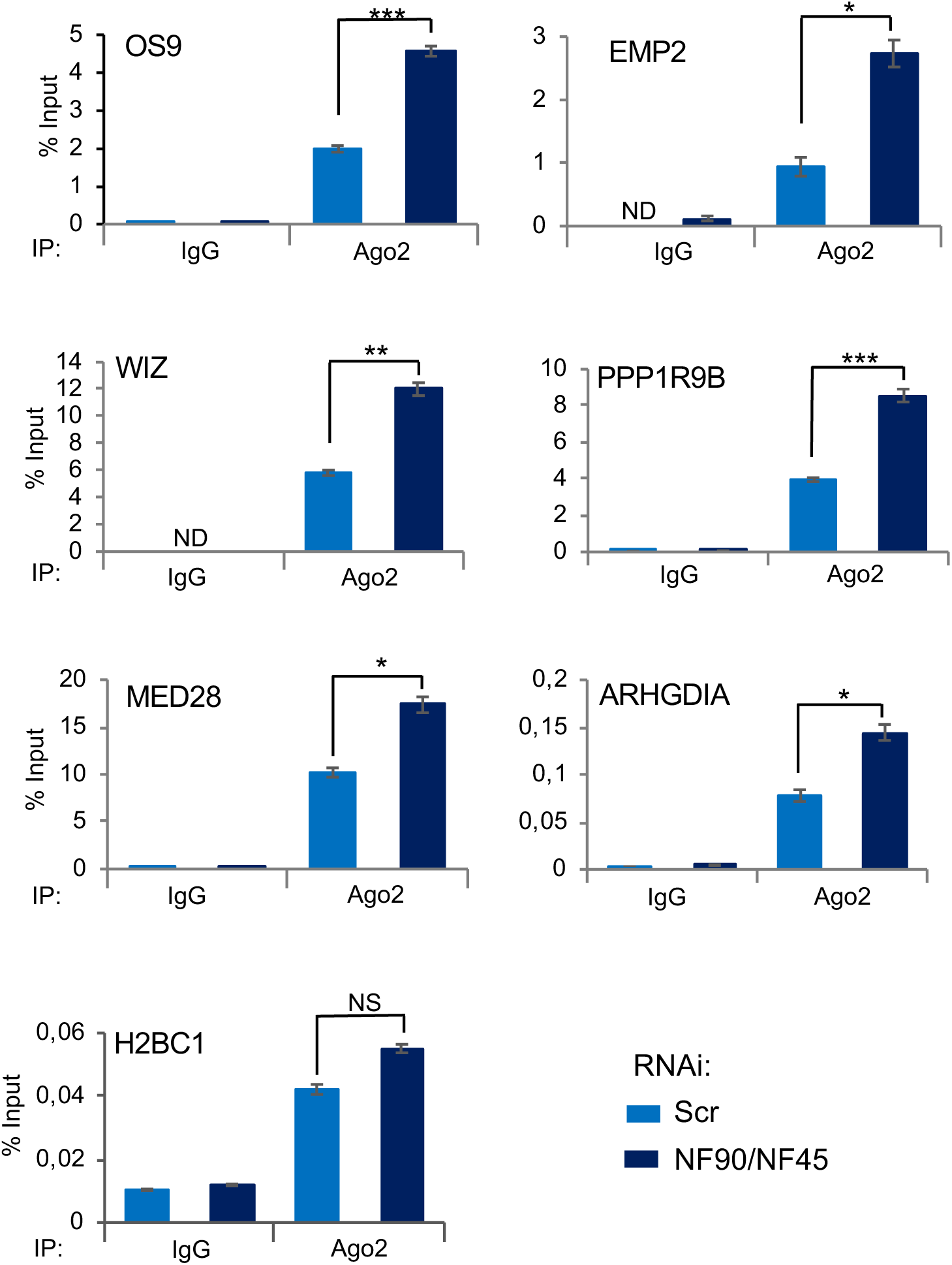
Downregulation of NF90/NF45 increases Ago2 binding to selected target mRNAs. RIP analysis of HEK293T cells transfected with NF90/NF45-targeting siRNAs or a non-targeting control (Scr), as indicated. RIPs were performed using anti-Ago2 or a control IgG antibody. Immunoprecipitates were analyzed using RT-qPCR. ND indicates ‘Not Detected’. Data represent mean ± SEM obtained from 3 independent experiments *(*P < 0.05, **P < 0.01, ***P < 0.001*, independent Student’s *t* test).

### NF90/NF45 impedes binding of Ago2 to VEGF mRNA during hypoxia

NF90 is known to stabilize VEGF mRNA during cancer-induced hypoxia [6]. We wondered whether the stabilization of VEGF by NF90 during hypoxia might be mediated by the ability of NF90 to influence Ago2 binding to VEGF mRNA. To this end, we performed RIP of Ago2 or IgG control after downregulation of NF90/NF45 and treatment with the hypoxia-inducing drug, CoCl_2_. HEK293T treated with 500 µM of CoCl_2_ show increased HIF1α expression (Figure 6A), confirming induction of hypoxia, and this concentration was used for further experiments. Depletion of NF90/NF45 had no effect on Ago2 expression in extracts or immunoprecipitates (Figure 6B, C). However, loss of NF90 significantly decreased the mRNA level of VEGF while increasing the level of H2BC1 mRNA (Figure 6D). RIP analysis showed that the binding of Ago2 to VEGF mRNA was significantly increased upon loss of NF90/NF45, while the binding to the negative control mRNA, H2BC1, did not significantly change (Figure 6E).

**Figure 6.**
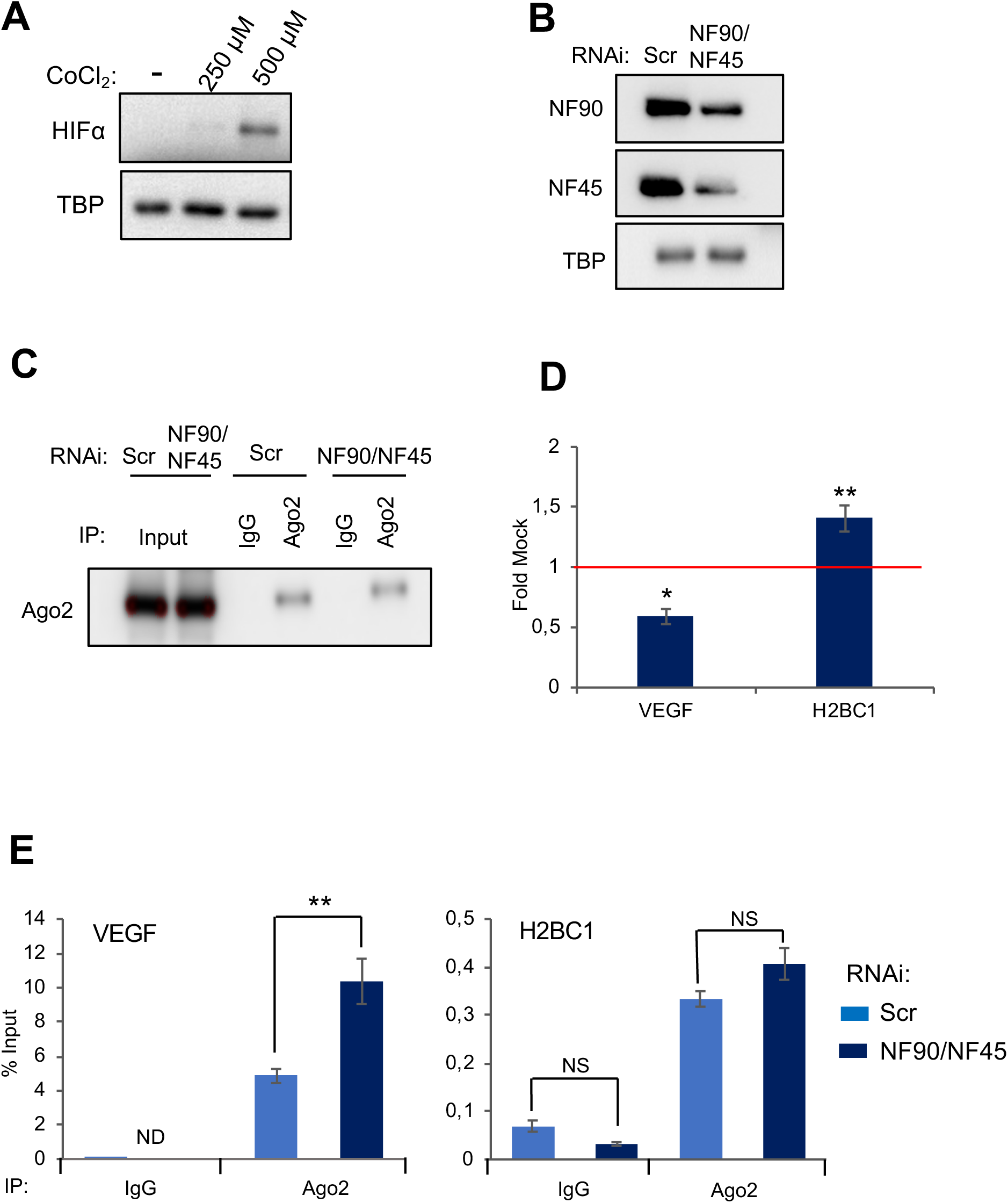
NF90/NF45 impedes binding of Ago2 to VEGF mRNA during hypoxia. **(A)** Extracts of HEK293T cells treated with different concentrations of CoCl_2_, as indicated, were analyzed by Western blot using antibodies to HIF1α and TBP. **(B)** Extracts of HEK293T cells transfected with siRNAs targeting NF90/NF45 or a non-targeting control (Scr) and treated with CoCl_2_ were analyzed by Western blot using the indicated antibodies. **(C) I**mmunoprecipitates obtained using anti-Ago2 or control IgG antibodies from extracts described in (B) were analyzed by Western blot using the indicated antibodies. **(D)** Total RNA obtained from HEK293T transfected with siRNAs targeting NF90 and NF45 or a non-targeting control (Scr) and treated with CoCl_2_ were analyzed by RT-qPCR. Data represent Fold Mock (IgG) relative to the control samples (siScr), which was attributed a value of 1 (red line), obtained from 3 independent experiments *(*P < 0.05, **P < 0.01, ***P < 0.001*, independent Student’s *t* test). **(E)** RIP analysis of HEK293T cells transfected with NF90/NF45-targeting siRNAs or a non-targeting control (Scr), as indicated, and treated with CoCl_2_. RIPs were performed using anti-Ago2 or a control IgG antibody. Immunoprecipitates were analyzed using RT-qPCR. ND indicates ‘Not Detected’. NS indicate ‘Not Significant’. Data represent mean ± SEM obtained from 3 independent experiments *(*P < 0.05, **P < 0.01, ***P < 0.001*, independent Student’s *t* test).

## DISCUSSION

The dsRNA binding protein, NF90, has been implicated in a number of cellular pathways, including the regulation of translation and RNA stability. However, the precise mechanisms involved in its different functions are not entirely clear. To better understand how NF90 performs these roles, we identified the interactome of cytoplasmic NF90. Identification of its partners may shed light on mechanisms by which NF90 is implicated in different pathways. Gene ontology analysis of the interactants revealed, not surprisingly, that almost all NF90 partners are involved in pathways in the processing, stability or translation of cellular RNA, consistent with the known functions of NF90. Interestingly, pathways associated with viral transcription and viral translation (Figure 1A) were among the significantly enriched gene ontology terms. This is notable since it has been shown that NF90 translocates from the nucleus to the cytoplasm as a consequence of viral infection and, following its nuclear export, NF90 was shown to bind viral mRNAs, playing a role in the antiviral immune response or enhancing viral replication, depending on the virus [2, 31]. We previously showed that nuclear NF90 is able to inhibit the biogenesis of several miRNAs involved in viral replication and antiviral response, such as miR-4753 and miR-3145 [13]. Consistently, these miRNAs were shown to be overexpressed in response to viral infection, inhibiting influenza A viral transcription and replication [32]. Thus, the identification of cytoplasmic NF90 partners involved in viral transcription and translation suggests that NF90 may play an important role in the cellular response to viral infection.

NF90, has been implicated in the regulation of translation and mRNA stability. Moreover, a recent study put forward the hypothesis that this might be due to its involvement in miRNA-mediated gene silencing [16]. However, a direct role for NF90 in RISC-mediated silencing had not so far been demonstrated. The interactome of cytoplasmic NF90 contained several components of RISC. These include DHX30, UPF1, DDX6 as well as the RISC-associated RNA helicase, MOV10. The effector of RISC-mediated silencing, Ago2, was also identified among NF90 interactants, which confirms previous reports identifying an interaction between NF90 and Ago2 [16, 23]. We furthermore determined that the association of NF90 with MOV10 and Ago2 occurs through RNA. NF90 likely exists in a cytosolic complex with RISC and RISC-associated factors since these factors co-sediment in a glycerol gradient. These data suggest that NF90 may be linked to RISC-mediated silencing via RNA-dependent interactions with RISC-associated proteins.

Analysis of NF90 and MOV10 CLIP data suggested that both proteins can associate with the 3’ UTR of a subset of target mRNAs. However, NF90 was shown to increase mRNA stability while MOV10 enhances RISC-mediated silencing. Since these effects appear contradictory, we wondered whether NF90 and MOV10 could compete for the binding of selected target mRNAs. Upon downregulation of MOV10, an increase in the association of NF90 to the target mRNAs was detected. Likewise, upon loss of the heterodimer NF90/NF45, an increase in MOV10 binding to the same mRNAs was measured. These findings suggest that NF90 and MOV10 might influence or interfere with the ability of the other factor to associate with its target RNAs. This interference is probably unlikely to occur through direct competition for the same binding sites, as reported for NF90 and Microprocessor binding to pri-miRNAs in the nucleus [3, 13, 33]. Indeed, the binding preferences for NF90 and MOV10 differ significantly. MOV10 binds ssRNA upstream of a structured region while NF90 appears to bind highly stable hairpin structures [13, 14]. The binding of one factor may modify RNA structure to disfavor binding of the other. Alternatively, one factor may recruit additional proteins that may influence the binding of the other factor. In any case, NF90 and MOV10 mutually interfere with each other’s binding to a subset of common target mRNAs.

MOV10 helicase activity is known to resolve mRNA structures in order to reveal obscured MREs within 3’ UTRs. This facilitates the binding of Ago2, which favors RISC-mediated silencing. Interestingly, depletion of NF90 significantly increased the association of Ago2 with target mRNAs. Consistent with increased Ago2 association, downregulation of the total level of the selected target mRNAs upon loss of NF90 was detected. These data suggest that NF90 enhances the stability of certain mRNAs by modulating the ability of Ago2 to associate with its target site and induce RISC-mediate silencing. It is interesting to note that NF90 has been identified as a subunit of P-bodies [34]. P-bodies are a site of mRNA storage as well as RISC-mediated mRNA degradation. It is tempting to speculate that NF90 within P-bodies may be implicated in the control of mRNA stability versus degradation by modulating the binding of RISC. It would also be interesting to determine whether the destabilization of mRNAs observed upon loss of NF90 occurs within P-bodies.

NF90 is known to translocate to the cytoplasm during cancer-induced hypoxia where it can bind and stabilize VEGF [5, 6]. Interestingly, we found that during hypoxia, loss of NF90/NF45 diminished of VEGF mRNA abundance, while its association with Ago2 was significantly increased. These results suggest that the role of NF90 in stabilizing VEGF during cancer-induced hypoxia might be the result of NF90 interfering with Ago2 binding to VEGF mRNA, and consequently, reduced targeting of VEGF mRNA by RISC activity. Therefore, better understanding of the role of NF90 in RISC-mediated silencing could potentially elucidate its effect on the fate of mRNAs involved in the antiviral immune response or during hypoxia induced in solid tumors.

## DECLARATIONS

### Availability of data and materials

All data generated or analysed during this study are included in this published article [and its supplementary information files].

### Authors’ contributions

GG performed experiments and contributed to writing the manuscript; CF helped with glycerol gradient sedimentation and RIP experiments; RNS helped with creating the stable cell line overexpressing NF90-FH and mass spectrometry; MB helped with RIP experiments; JB created the stable cell line overexpressing NF90-FH and prepared samples for mass spectrometry; RK designed the study and wrote the manuscript. All authors read and approved the final manuscript.

## Acknowledgements

The authors thank Claudio Lorenzi for bioinformatic assistance, Thomas Lecesne for technical assistance and members of the Gene Regulation lab for helpful discussions.

## REFERENCES

1. Castella S, Bernard R, Corno M, Fradin A, Larcher JC: Ilf3 and NF90 functions in RNA biology. Wiley Interdiscip Rev RNA 2014.

2. Li X, Liu CX, Xue W, Zhang Y, Jiang S, Yin QF, Wei J, Yao RW, Yang L, Chen LL: Coordinated circRNA Biogenesis and Function with NF90/NF110 in Viral Infection. Mol Cell 2017, 67(2):214–227 e217.

3. Sakamoto S, Aoki K, Higuchi T, Todaka H, Morisawa K, Tamaki N, Hatano E, Fukushima A, Taniguchi T, Agata Y: The NF90-NF45 complex functions as a negative regulator in the microRNA processing pathway. Mol Cell Biol 2009, 29(13):3754–3769.

4. Harashima A, Guettouche T, Barber GN: Phosphorylation of the NFAR proteins by the dsRNA-dependent protein kinase PKR constitutes a novel mechanism of translational regulation and cellular defense. Genes Dev 2010, 24(23):2640–2653.

5. Vrakas CN, Herman AB, Ray M, Kelemen SE, Scalia R, Autieri MV: RNA stability protein ILF3 mediates cytokine-induced angiogenesis. FASEB J 2019, 33(3):3304–3316.

6. Zhang W, Xiong Z, Wei T, Li Q, Tan Y, Ling L, Feng X: Nuclear factor 90 promotes angiogenesis by regulating HIF-1alpha/VEGF-A expression through the PI3K/Akt signaling pathway in human cervical cancer. Cell Death Dis 2018, 9(3):276.

7. Patino C, Haenni AL, Urcuqui-Inchima S: NF90 isoforms, a new family of cellular proteins involved in viral replication? Biochimie 2015, 108:20–24.

8. Song D, Huang H, Wang J, Zhao Y, Hu X, He F, Yu L, Wu J: NF90 regulates PARP1 mRNA stability in hepatocellular carcinoma. Biochem Biophys Res Commun 2017, 488(1):211–217.

9. Vumbaca F, Phoenix KN, Rodriguez-Pinto D, Han DK, Claffey KP: Double-stranded RNA-binding protein regulates vascular endothelial growth factor mRNA stability, translation, and breast cancer angiogenesis. Mol Cell Biol 2008, 28(2):772–783.

10. Nourreddine S, Lavoie G, Paradis J, Ben El Kadhi K, Meant A, Aubert L, Grondin B, Gendron P, Chabot B, Bouvier M et al: NF45 and NF90 Regulate Mitotic Gene Expression by Competing with Staufen-Mediated mRNA Decay. Cell Rep 2020, 31(7):107660.

11. Jayachandran U, Grey H, Cook AG: Nuclear factor 90 uses an ADAR2-like binding mode to recognize specific bases in dsRNA. Nucleic Acids Res 2016, 44(4):1924–1936.

12. Schmidt T, Knick P, Lilie H, Friedrich S, Golbik RP, Behrens SE: Coordinated Action of Two Double-Stranded RNA Binding Motifs and an RGG Motif Enables Nuclear Factor 90 To Flexibly Target Different RNA Substrates. Biochemistry 2016, 55(6):948–959.

13. Grasso G, Higuchi T, Mac V, Barbier J, Helsmoortel M, Lorenzi C, Sanchez G, Bello M, Ritchie W, Sakamoto S et al: NF90 modulates processing of a subset of human pri-miRNAs. Nucleic Acids Res 2020, 48(12):6874–6888.

14. Gwizdek C, Ossareh-Nazari B, Brownawell AM, Evers S, Macara IG, Dargemont C: Minihelix-containing RNAs mediate exportin-5-dependent nuclear export of the double-stranded RNA-binding protein ILF3. J Biol Chem 2004, 279(2):884–891.

15. Schmidt T, Knick P, Lilie H, Friedrich S, Golbik RP, Behrens SE: The properties of the RNA-binding protein NF90 are considerably modulated by complex formation with NF45. Biochem J 2017, 474(2):259–280.

16. Idda ML, Lodde V, McClusky WG, Martindale JL, Yang X, Munk R, Steri M, Orru V, Mulas A, Cucca F et al: Cooperative translational control of polymorphic BAFF by NF90 and miR-15a. Nucleic Acids Res 2018, 46(22):12040–12051.

17. Flores O, Kennedy EM, Skalsky RL, Cullen BR: Differential RISC association of endogenous human microRNAs predicts their inhibitory potential. Nucleic Acids Res 2014, 42(7):4629–4239.

18. Bartel DP: MicroRNAs: target recognition and regulatory functions. Cell 2009, 136(2):215–233.

19. Hock J, Weinmann L, Ender C, Rudel S, Kremmer E, Raabe M, Urlaub H, Meister G: Proteomic and functional analysis of Argonaute-containing mRNA-protein complexes in human cells. EMBO Rep 2007, 8(11):1052–1060.

20. Gregersen LH, Schueler M, Munschauer M, Mastrobuoni G, Chen W, Kempa S, Dieterich C, Landthaler M: MOV10 Is a 5’ to 3’ RNA helicase contributing to UPF1 mRNA target degradation by translocation along 3’ UTRs. Mol Cell 2014, 54(4):573–585.

21. Huang F, Zhang J, Zhang Y, Geng G, Liang J, Li Y, Chen J, Liu C, Zhang H: RNA helicase MOV10 functions as a co-factor of HIV-1 Rev to facilitate Rev/RRE-dependent nuclear export of viral mRNAs. Virology 2015, 486:15–26.

22. Liu T, Sun Q, Liu Y, Cen S, Zhang Q: The MOV10 helicase restricts hepatitis B virus replication by inhibiting viral reverse transcription. J Biol Chem 2019, 294(51):19804–19813.

23. Meister G, Landthaler M, Peters L, Chen PY, Urlaub H, Luhrmann R, Tuschl T: Identification of novel argonaute-associated proteins. Curr Biol 2005, 15(23):2149–2155.

24. Yang Q, Jankowsky E: The DEAD-box protein Ded1 unwinds RNA duplexes by a mode distinct from translocating helicases. Nat Struct Mol Biol 2006, 13(11):981–986.

25. Kenny PJ, Kim M, Skariah G, Nielsen J, Lannom MC, Ceman S: The FMRP-MOV10 complex: a translational regulatory switch modulated by G-Quadruplexes. Nucleic Acids Res 2020, 48(2):862–878.

26. Kenny PJ, Zhou H, Kim M, Skariah G, Khetani RS, Drnevich J, Arcila ML, Kosik KS, Ceman S: MOV10 and FMRP regulate AGO2 association with microRNA recognition elements. Cell Rep 2014, 9(5):1729–1741.

27. Contreras X, Salifou K, Sanchez G, Helsmoortel M, Beyne E, Bluy L, Pelletier S, Rousset E, Rouquier S, Kiernan R: Nuclear RNA surveillance complexes silence HIV-1 transcription. PLoS Pathog 2018, 14(3):e1006950.

28. Nussbacher JK, Yeo GW: Systematic Discovery of RNA Binding Proteins that Regulate MicroRNA Levels. Mol Cell 2018, 69(6):1005–1016 e1007.

29. Guan D, Altan-Bonnet N, Parrott AM, Arrigo CJ, Li Q, Khaleduzzaman M, Li H, Lee CG, Pe’ery T, Mathews MB: Nuclear factor 45 (NF45) is a regulatory subunit of complexes with NF90/110 involved in mitotic control. Mol Cell Biol 2008, 28(14):4629–4641.

30. Wolkowicz UM, Cook AG: NF45 dimerizes with NF90, Zfr and SPNR via a conserved domain that has a nucleotidyltransferase fold. Nucleic Acids Res 2012, 40(18):9356–9368.

31. Isken O, Baroth M, Grassmann CW, Weinlich S, Ostareck DH, Ostareck-Lederer A, Behrens SE: Nuclear factors are involved in hepatitis C virus RNA replication. RNA 2007, 13(10):1675–1692.

32. Khongnomnan K, Makkoch J, Poomipak W, Poovorawan Y, Payungporn S: Human miR-3145 inhibits influenza A viruses replication by targeting and silencing viral PB1 gene. Exp Biol Med (Maywood) 2015, 240(12):1630–1639.

33. Barbier J, Chen X, Sanchez G, Cai M, Helsmoortel M, Higuchi T, Giraud P, Contreras X, Yuan G, Feng Z et al: An NF90/NF110-mediated feedback amplification loop regulates dicer expression and controls ovarian carcinoma progression. Cell Res 2018, 28(5):556–571.

34. Hubstenberger A, Courel M, Benard M, Souquere S, Ernoult-Lange M, Chouaib R, Yi Z, Morlot JB, Munier A, Fradet M et al: P-Body Purification Reveals the Condensation of Repressed mRNA Regulons. Mol Cell 2017, 68(1):144–157 e145.

