## Supplementary material for "NF90 Interacts with Components of RISC and Modulates Association of Ago2 with mRNA": supplemntal figures

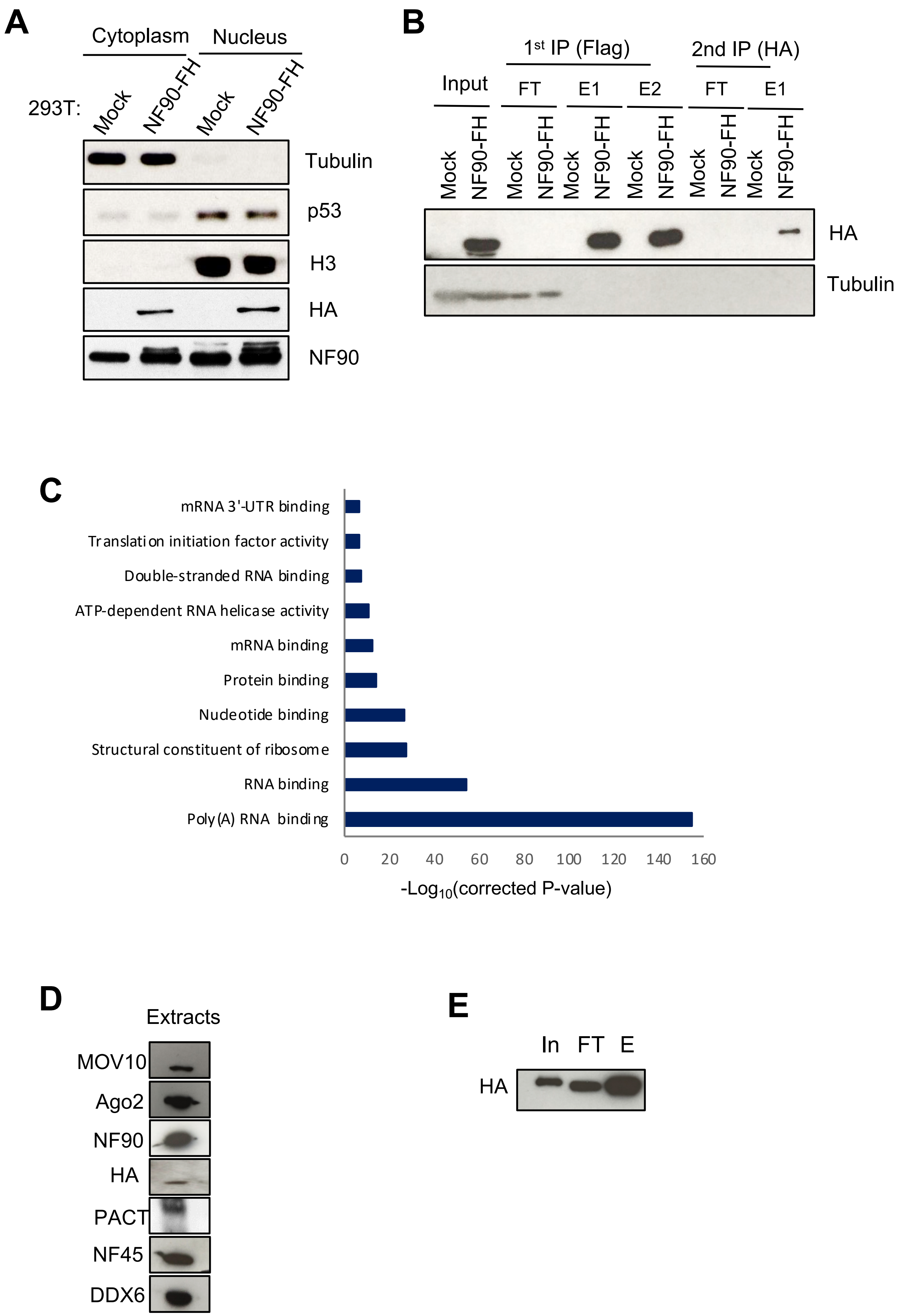
A
Cytoplasm
Nucleus
293T:
Mock
NF90-FH
Mock
NF90-FH
Tubulin
p53
H3
HA
NF90
B
1st IP (Flag)
2nd IP (HA)
Input
FT
E1
E2
FT
E1
Mock
NF90-FH
HA
Tubulin
C
mRNA 3'-UTR binding
Translation initiation factor activity
Double-stranded RNA binding
ATP-dependent RNA helicase activity
mRNA binding
Protein binding
Nucleotide binding
Structural constituent of ribosome
RNA binding
Poly(A) RNA binding
0
20
40
60
80
100
120
140
160
-Log10(corrected P-value)
D
Extracts
MOV10
Ago2
NF90
HA
PACT
NF45
DDX6
E
In
FT
E
HA

**Figure S1**

**Figure S1.** NF90 interacts with proteins involved in translational repression and RNA processing. **(A)** Cytoplasmic and nuclear extracts of WT (mock) and NF90-FH stably overexpressing (NF90-FH) HEK293T cells were analyzed by Western blot using the indicated antibodies. **(B)** Cytoplasmic extracts described in A underwent tandem affinity purification using FLAG and HA antibodies. Samples were analyzed by western blot using the indicated antibodies (FT= flow through, E1=elution 1; E2=elution 2). **(C)** Molecular functions of NF90-associated proteins identified by mass spectrometry were analyzed using gene ontology. **(D) C**ytoplasmic extracts of NF90-FH overexpressing HEK293T cells used for FLAG immunoprecipitation followed by glycerol gradient sedimentations were analyzed by western blot using the antibodies indicated. **(E)** An aliquot of FLAG immunoprecipitate from NF90-FH overexpressing HEK293T cells used for glycerol gradient sedimentation were analyzed by western blot, using the antibodies indicated (In= input, FT= flowthrough, E= elution) .

### Figure S1

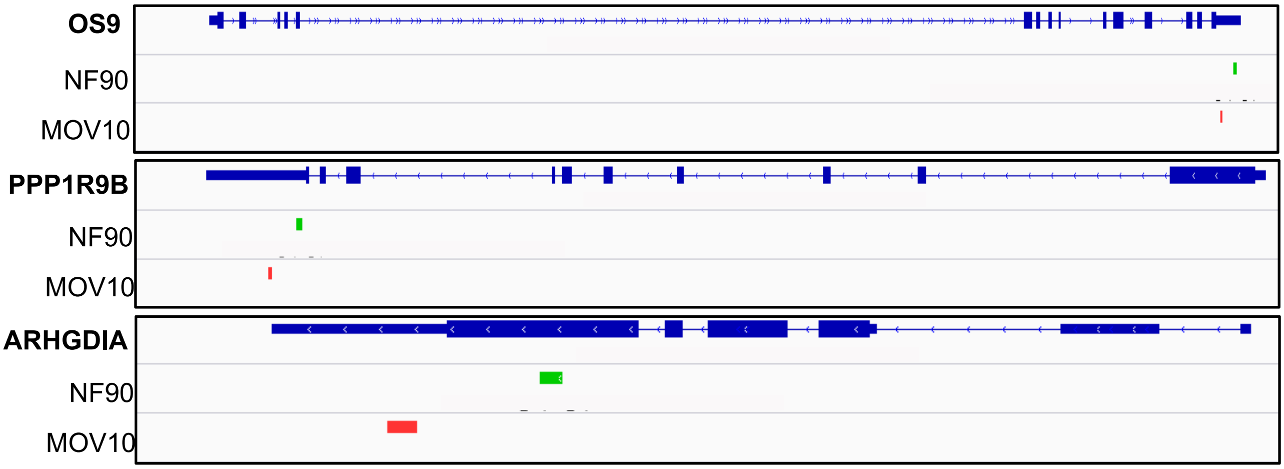


**Figure S2.** NF90 and MOV10 can bind the same target mRNAs. Screenshots of NF90 and MOV10 eCLIP and iCLIP, respectively, showing regions associated with NF90 (green bars) and MOV10 (red bars) within selected target mRNAs. Introns are shown as thin lines, 5’ and 3’ UTRs are shown as medium lines and exons are shown as thick lines. The orientation of the transcript with respect to the genome is indicated by arrows.

**Figure S2**

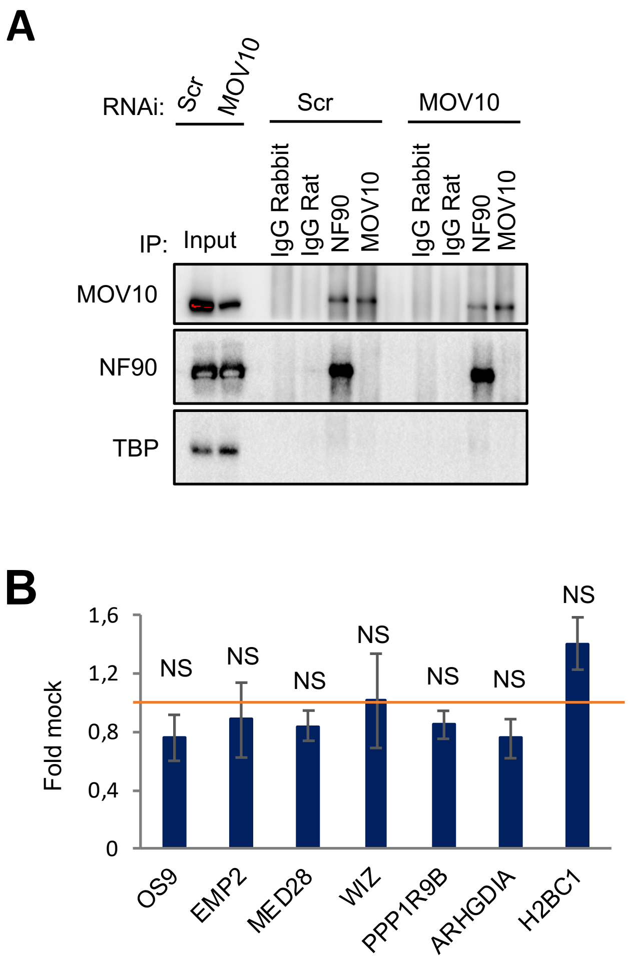


**Figure S3.** MOV10 modulates NF90 association with common target mRNAs. **(A)** Extracts of HEK293T cells transfected with siRNAs targeting MOV10 or a non-targeting control (Scr) and immunoprecipitates obtained using antibodies anti-NF90, anti-MOV10 or control IgG were analyzed by Western blot using the indicated antibodies. **(B)** Total RNA obtained from HEK293T transfected with siRNAs targeting MOV10 or a non-targeting control (Scr) was analyzed by RT-qPCR. NS indicates 'Not Significant'. Data represent Fold Mock (IgG) relative to the control samples (siScr), which was attributed a value of 1 (red line), obtained from 4 independent experiments (*$^*P < 0.05$, $^{**}P < 0.01$, $^{***}P < 0.001$*, independent Student's *t* test).

**Figure S3**

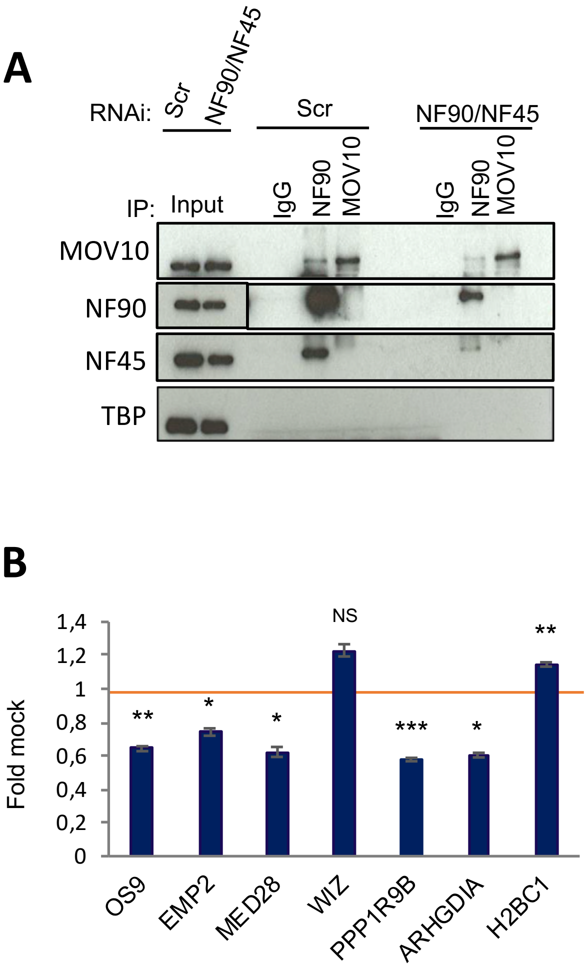


**Figure S4.** NF90 modulates MOV10 association with common target mRNAs. **(A)** Extracts of HEK293T cells transfected with siRNAs targeting NF90 and NF45 or a non-targeting control (Scr) and immunoprecipitates obtained using antibodies anti-NF90, anti-MOV10 or control IgG were analyzed by Western blot using the indicated antibodies. **(B)** Total RNA obtained from HEK293T transfected with siRNAs targeting NF90 and NF45 or a non-targeting control (Scr) was analyzed by RT-qPCR. NS indicates 'Not Significant'. Data represent Fold Mock (IgG) relative to the control samples (siScr), which was attributed a value of 1 (red line), obtained from 4 independent experiments *(*P < 0.05, **P < 0.01, ***P < 0.001*, independent Student's *t* test).

**Figure S4**

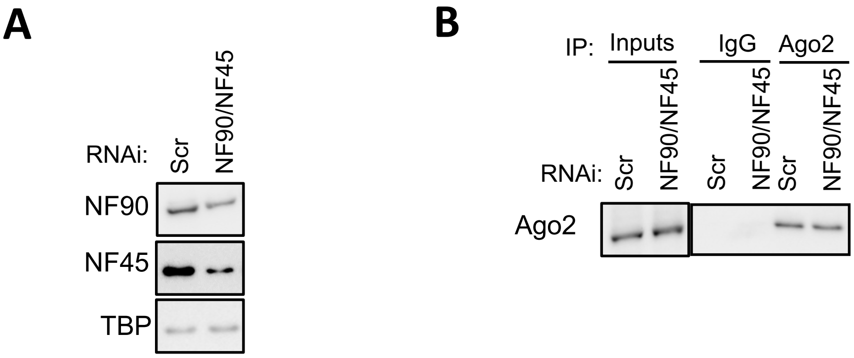


**Figure S5.** Downregulation of NF90/NF45 does not affect the expression of Ago2 **(A)** Extracts of HEK293T cells transfected with siRNAs targeting NF90/NF45 or a non-targeting control (Scr) were analyzed by Western blot using the indicated antibodies. **(B) I**mmunoprecipitates obtained using anti-Ago2 or control IgG antibodies from extracts of HEK293T cells transfected with siRNAs targeting NF90/NF45 or a non-targeting control (Scr) were analyzed by Western blot using anti-Ago2 antibody.

### Figure S5

**Supplementary Table S1.** Double stranded siRNAs used in this study.

| siRNA | Sequence (5' to 3') |
|---|---|
| Scr | gcgcgcuuuguaggauucg(dTdT) |
| NF45 | guggugauacucaagauucugccaa(dTdT) |
| NF90 | ccaaggaacucuaucacaa(dTdT) |
| MOV10 | guggaauuggaccgugucaagcuga(dTdT) |

**Supplementary Table S2.** Primary antibodies used in this study.

| Antibody | Reference | Supplier |
|---|---|---|
| NF90 | A303-651A | Bethyl Laboratories |
| NF45 | A303-147A | Bethyl Laboratories |
| MOV10 | A301-571A | Bethyl Laboratories |
| PACT | sc-81569 | SCBT |
| HA | 12CA5 | Roche |
| DDX6 | A300-461 | Bethyl Laboratories |
| Ago2 WB | MABE253, clone 119A | Sigma-Aldrich |
| Ago2 RIP | 03-110 | Sigma-Aldrich |
| TUBULIN | DM1A clone, T6199 | Sigma-Aldrich |
| TBP | sc-421 | SCBT |

**Supplementary Table S3.** Primers used in this study.

| Primer | Forward (5' to 3') | Reverse (5' to 3') |
|---|---|---|
| OS9 | CTGCCCTTAGTGATGTTTGGA | ATTCTACCATTTCCTCAGCAACA |
| EMP2 | GACCCATCCACCATTCATTC | CCACTGTACCCAGCCAGTTT |
| MED28 | GAGCAGGCAGTAGGATGAGG | CCCCTCCACAGGGATATTTT |
| WIZ | TTGGCTGCTCCTTCTTGTTT | GCCTGTTAATCCCTCCTTCC |
| PPP1R9B | GCCCTGCACTGATTTCTCAT | TATTTGGCACCTGGAAGAGG |
| ARHGDIA | TGCCTCTGCCTTTTCTGTCT | GCACTTGGTCCCTTGTTTGT |
| H2BC1 | GTGCTAAAGCAGGTCCATCC | GCATGTTTAGCCAGCTCTCC |
| VEGFA | TGACAGGGAAGAGGAGGAGA | CGTCTGACCTGGGGTAGAGA |

**Supplementary Table S4.** Proteins associated with NF90 in the cytoplasm, identified by mass spectrometry.

| Reference | Gene Symbol |
|---|---|
| Q99848_EBP2_HUMAN | EBNA1BP2 |
| Q13595_TRA2A_HUMAN | TRA2A |
| P16403_H12_HUMAN | HIST1H1C |
| P82650_RT22_HUMAN | MRPS22 |
| O75175_CNOT3_HUMAN | CNOT3 |
| Q13573_SNW1_HUMAN | SNW1 |
| P78344_IF4G2_HUMAN | EIF4G2 |
| Q96KR1_ZFR_HUMAN | ZFR |
| Q96PU8_QKI_HUMAN | QKI |
| P42696_RBM34_HUMAN | RBM34 |
| Q96QR8_PURB_HUMAN | PURB |
| Q9NX24_NHP2_HUMAN | NHP2 |
| Q9NX05_F120C_HUMAN | FAM120C |
| Q9ULX6_AKP8L_HUMAN | AKAP8L |
| Q99575_POP1_HUMAN | POP1 |
| Q9BYJ9_YTHD1_HUMAN | YTHDF1 |
| P16383_GCFC2_HUMAN | GCFC2 |
| Q8IX01_SUGP2_HUMAN | SUGP2 |
| Q9UL18_AGO1_HUMAN | AGO1 |
| Q8IZH2_XRN1_HUMAN | XRN1 |
| Q9Y265_RUVB1_HUMAN | RUVBL1 |
| P33993_MCM7_HUMAN | MCM7 |
| Q9UKV3_ACINU_HUMAN | ACIN1 |
| O75822_EIF3J_HUMAN | EIF3J |
| Q9H307_PININ_HUMAN | PNN |
| P42357_HUTH_HUMAN | HAL |
| O43242_PSMD3_HUMAN | PSMD3 |
| P13639_EF2_HUMAN | EEF2 |
| P35251_RFC1_HUMAN | RFC1 |
| P08621_RU17_HUMAN | SNRNP70 |
| P0CB38_PAB4L_HUMAN | PABPC4L |
| Q02539_H11_HUMAN | HIST1H1A |
| P42677_RS27_HUMAN | RPS27 |
| Q5JNZ5_RS26L_HUMAN | RPS26P11 |
| Q16695_H31T_HUMAN | HIST3H3 |
| P13995_MTDC_HUMAN | MTHFD2 |
| P05386_RLA1_HUMAN | RPLP1 |
| Q01804_OTUD4_HUMAN | OTUD4 |
| Q6EMK4_VASN_HUMAN | VASN |
| O43709_WBS22_HUMAN | WBSCR22 |
| O43347_MSI1H_HUMAN | MSI1 |
| Q15365_PCBP1_HUMAN | PCBP1 |
| Q15287_RNPS1_HUMAN | RNPS1 |
| P61513_RL37A_HUMAN | RPL37A |
| Q7Z2T5_TRM1L_HUMAN | TRMT1L |
| P09012_SNRPA_HUMAN | SNRPA |
| P29558_RBMS1_HUMAN | RBMS1 |
| Q9BRZ2_TRI56_HUMAN | TRIM56 |
| P19525_E2AK2_HUMAN | EIF2AK2 |
| O75934_SPF27_HUMAN | BCAS2 |
| P55209_NP1L1_HUMAN | NAP1L1 |
| P62314_SMD1_HUMAN | SNRPD1 |
| Q3MHD2_LSM12_HUMAN | LSM12 |
| Q6ZN17_LN28B_HUMAN | LIN28B |
| Q58A45_PAN3_HUMAN | PAN3 |
| Q9ULR0_ISY1_HUMAN | ISY1 |
| Q8N4Q0_ZADH2_HUMAN | ZADH2 |
| Q14498_RBM39_HUMAN | RBM39 |
| Q07666_KHDR1_HUMAN | KHDRBS1 |
| P24534_EF1B_HUMAN | EEF1B2 |
| Q9Y383_LC7L2_HUMAN | LUC7L2 |
| P50991_TCPD_HUMAN | CCT4 |
| P08238_HS90B_HUMAN | HSP90AB1 |
| O75494_SRS10_HUMAN | SRSF10 |
| P60709_ACTB_HUMAN | ACTB |
| Q15366_PCBP2_HUMAN | PCBP2 |
| P04844_RPN2_HUMAN | RPN2 |
| P25705_ATPA_HUMAN | ATP5A1 |
| P60891_PRPS1_HUMAN | PRPS1 |
| Q9BYD3_RM04_HUMAN | MRPL4 |
| O43167_ZBT24_HUMAN | ZBTB24 |
| Q9UHB9_SRP68_HUMAN | SRP68 |
| K7ER90_K7ER90_HUMAN | EIF3G |
| Q9BUF5_TBB6_HUMAN | TUBB6 |
| P11908_PRPS2_HUMAN | PRPS2 |
| Q9NQ92_COPRS_HUMAN | COPRS |
| P32119_PRDX2_HUMAN | PRDX2 |
| Q02809_PLOD1_HUMAN | PLOD1 |
| Q5BKZ1_ZN326_HUMAN | ZNF326 |

| Reference | Gene Symbol |
|---|---|
| Q15434_RBMS2_HUMAN | RBMS2 |
| Q14558_KPRA_HUMAN | PRPSAP1 |
| O14818_PSA7_HUMAN | PSMA7 |
| Q58FF8_H90B2_HUMAN | HSP90AB2P |
| O43734_CIKS_HUMAN | TRAF3IP2 |
| Q9BWU0_NADAP_HUMAN | SLC4A1AP |
| Q96EC8_YIPF6_HUMAN | YIPF6 |
| P20618_PSB1_HUMAN | PSMB1 |
| P98179_RBM3_HUMAN | RBM3 |
| Q13243_SRSF5_HUMAN | SRSF5 |
| Q8N5C8_TAB3_HUMAN | TAB3 |
| Q5VYS8_TUT7_HUMAN | ZCCHC6 |
| J3QR62_J3QR62_HUMAN | DDX5 |
| P05141_ADT2_HUMAN | SLC25A5 |
| Q07955_SRSF1_HUMAN | SRSF1 |
| Q14671_PUM1_HUMAN | PUM1 |
| Q9NZN8_CNOT2_HUMAN | CNOT2 |
| B7Z645_B7Z645_HUMAN | SYNCRIP |
| Q15084_PDIA6_HUMAN | PDIA6 |
| P62273_RS29_HUMAN | RPS29 |
| Q9NP73_ALG13_HUMAN | ALG13 |
| Q9NY12_GAR1_HUMAN | GAR1 |
| O00567_NOP56_HUMAN | NOP56 |
| Q92600_RCD1_HUMAN | RQCD1 |
| Q9BTZ2_DHRS4_HUMAN | DHRS4 |
| P49327_FAS_HUMAN | FASN |
| Q14694_UBP10_HUMAN | USP10 |
| Q16629_SRSF7_HUMAN | SRSF7 |
| C9JUF0_C9JUF0_HUMAN | EIF4A2 |
| Q9BZI7_REN3B_HUMAN | UPF3B |
| Q9HCS7_SYF1_HUMAN | XAB2 |
| Q9H7E9_CH033_HUMAN | C8orf33 |
| Q9NWU5_RM22_HUMAN | MRPL22 |
| Q9Y3D9_RT23_HUMAN | MRPS23 |
| P26368_U2AF2_HUMAN | U2AF2 |
| Q96HS1_PGAM5_HUMAN | PGAM5 |
| O75940_SPF30_HUMAN | SMNDC1 |
| P42704_LPPRC_HUMAN | LRPPRC |
| O43172_PRP4_HUMAN | PRPF4 |
| C9JQR9_C9JQR9_HUMAN | RPSAP58 |
| Q96L21_RL10L_HUMAN | RPL10L |

| Reference | Gene Symbol |
|---|---|
| Q08378_GOGA3_HUMAN | GOLGA3 |
| Q3KQU3_MA7D1_HUMAN | MAP7D1 |
| E9PRG8_CK098_HUMAN | C11orf98 |
| O14980_XPO1_HUMAN | XPO1 |
| Q5LJB1_Q5LJB1_HUMAN | UCHL5 |
| P49736_MCM2_HUMAN | MCM2 |
| Q8N8E3_CE112_HUMAN | CEP112 |
| O75152_ZC11A_HUMAN | ZC3H11A |
| Q8WXX5_DNJC9_HUMAN | DNAJC9 |
| Q9NVU7_SDA1_HUMAN | SDAD1 |
| O00505_IMA4_HUMAN | KPNA3 |
| O00458_IFRD1_HUMAN | IFRD1 |
| P51659_DHB4_HUMAN | HSD17B4 |
| Q96PX6_CC85A_HUMAN | CCDC85A |
| P08579_RU2B_HUMAN | SNRPB2 |
| Q5QJ74_TBCEL_HUMAN | TBCEL |
| Q9UKV8_AGO2_HUMAN | AGO2 |
| O15294_OGT1_HUMAN | OGT |
| P62807_H2B1C_HUMAN | HIST1H2BC |
| Q9BQ39_DDX50_HUMAN | DDX50 |
| O75962_TRIO_HUMAN | TRIO |
| P25789_PSA4_HUMAN | PSMA4 |
| O95721_SNP29_HUMAN | SNAP29 |
| Q9NRW3_ABC3C_HUMAN | APOBEC3C |
| O60437_PEPL_HUMAN | PPL |
| P30872_SSR1_HUMAN | SSTR1 |
| Q5TZA2_CROCC_HUMAN | CROCC |
